# Molecular differences between neonatal and adult stria vascularis from organotypic explants and transcriptomics

**DOI:** 10.1101/2024.04.24.590986

**Authors:** Matsya Ruppari Thulasiram, Ryosuke Yamamoto, Rafal T. Olszewski, Shoujun Gu, Robert J. Morell, Michael Hoa, Alain Dabdoub

## Abstract

**Summary:** The stria vascularis (SV), part of the blood-labyrinth barrier, is an essential component of the inner ear that regulates the ionic environment required for hearing. SV degeneration disrupts cochlear homeostasis, leading to irreversible hearing loss, yet a comprehensive understanding of the SV, and consequently therapeutic availability for SV degeneration, is lacking. We developed a whole-tissue explant model from neonatal and adult mice to create a robust platform for SV research. We validated our model by demonstrating that the proliferative behaviour of the SV *in vitro* mimics SV *in vivo,* providing a representative model and advancing high-throughput SV research. We also provided evidence for pharmacological intervention in our system by investigating the role of Wnt/β-catenin signaling in SV proliferation. Finally, we performed single-cell RNA sequencing from *in vivo* neonatal and adult mouse SV and revealed key genes and pathways that may play a role in SV proliferation and maintenance. Together, our results contribute new insights into investigating biological solutions for SV-associated hearing loss.

**Significance:** Hearing loss impairs our ability to communicate with people and interact with our environment. This can lead to social isolation, depression, cognitive deficits, and dementia. Inner ear degeneration is a primary cause of hearing loss, and our study provides an in depth look at one of the major sites of inner ear degeneration: the stria vascularis. The stria vascularis and associated blood-labyrinth barrier maintain the functional integrity of the auditory system, yet it is relatively understudied. By developing a new *in vitro* model for the young and adult stria vascularis and using single cell RNA sequencing, our study provides a novel approach to studying this tissue, contributing new insights and widespread implications for auditory neuroscience and regenerative medicine.

**Highlights:**

- We established an organotypic explant system of the neonatal and adult stria vascularis with an intact blood-labyrinth barrier.
- Proliferation of the stria vascularis decreases with age *in vitro*, modelling its proliferative behaviour *in vivo*.
- Pharmacological studies using our *in vitro* SV model open possibilities for testing injury paradigms and therapeutic interventions.
- Inhibition of Wnt signalling decreases proliferation in neonatal stria vascularis.
- We identified key genes and transcription factors unique to developing and mature SV cell types using single cell RNA sequencing.

## INTRODUCTION

Hearing relies on the regulation of cochlear homeostasis by the stria vascularis (SV). The SV lines the lateral wall of the inner ear and is comprised of three epithelial cell layers: the marginal layer originates from the otic epithelium and faces the endolymph of the scala media (Kikuchi & Hilding, 1966); the basal cells arise from the otic mesenchyme and are connected to the spiral ligament fibrocytes (Liu et al., 2017; Locher et al., 2005; Trowe et al., 2011); and the intermediate cell layer is made up of specialized perivascular resident macrophage-like melanocytes (ie. intermediate cells) derived from the neural crest (Morell et al., 2020; Renauld et al., 2022; Shibata et al., 2016; Steel & Barkway, 1989; Udagawa et al., 2024) and a network of endothelial cells and pericytes that make up the blood-labyrinth barrier (BLB; (Shi, 2011; Carraro et al., 2016; Figure 1A-B). The SV generates and maintains the endocochlear potential that is required for sound transduction, and protects the inner ear against pathogenic infiltration, making it crucial for hearing. SV degeneration can occur due to aging, ototoxic drugs, and genetic disease. Once damaged, the SV does not have the capacity to regenerate, which leads to progressive and irreversible hearing loss (reviewed in Thulasiram et al., 2022). Therefore, there is a need to better understand the SV and develop regenerative therapies targeting the SV.

**Figure 1.**
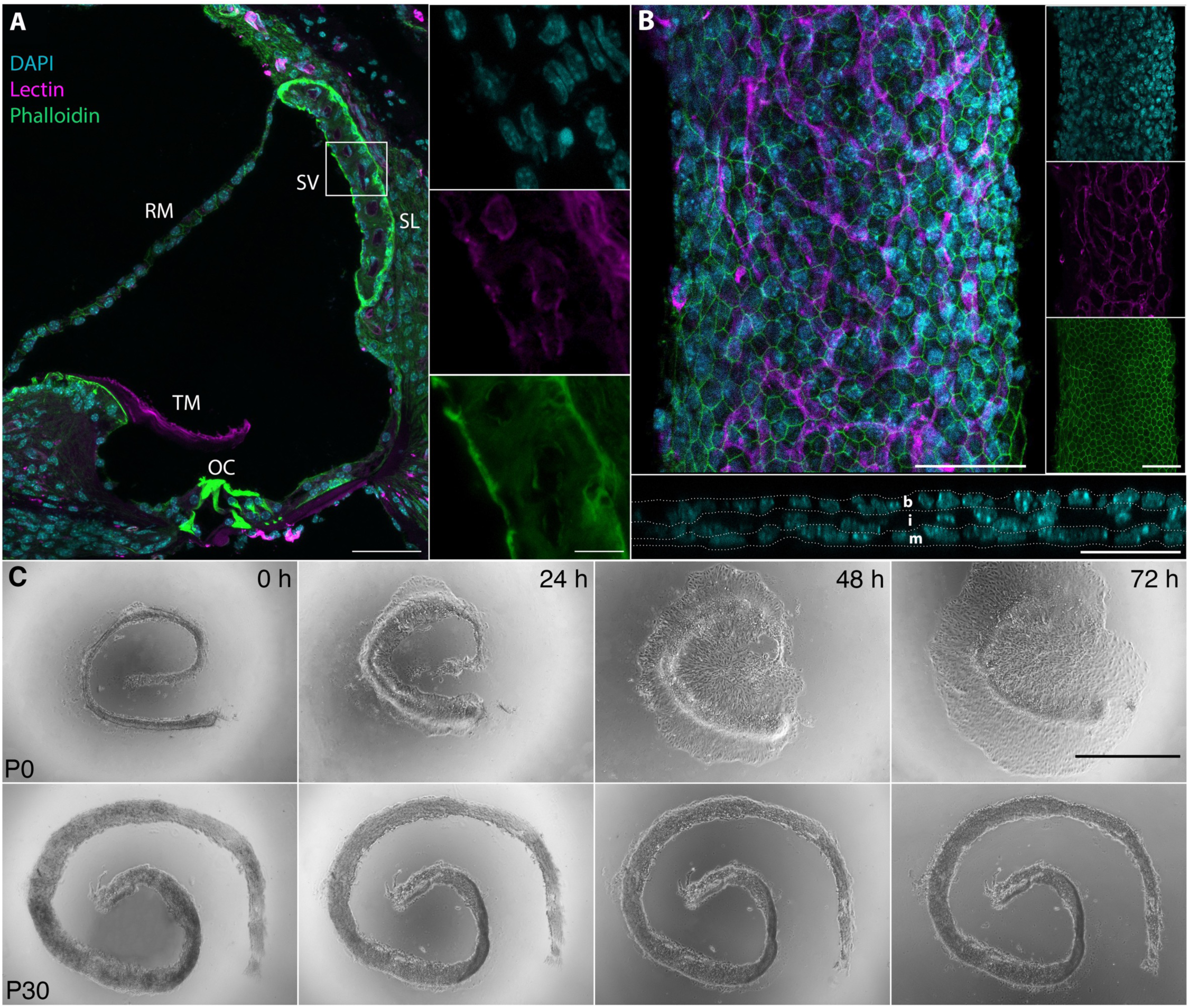
Generating an *in vitro* organotypic explant model for the neonatal and adult mouse stria vascularis. P30 cross section of a cochlear turn (A) and P0 wholemount stria vascularis (B) stained with Phalloidin for actin (green) Lectin for blood vessels (magenta) and DAPI for nuclei (blue). Orthogonal view of the wholemount SV in bottom panel of (B) marks the nuclei of the three cell layers of the SV (white dashed lines; m= marginal cells, i = intermediate cells, b = basal cells). Scale bars in A: 50 µm in low magnification images, 10 µm in high magnification images. Scale bars in B: 50 µm. C) Side-by-side differential interference contrast imaging of neonatal and adult SV organotypic explants. SV were dissected and cultured on Matrigel-coated plates in standard conditions for 72 h. Scale bar: 1 mm.

One of the main limitations for advancing SV research is the lack of an *in vitro* system that investigates the SV as a whole tissue. Techniques that currently exist include isolation, purification, and culture of the individual SV or BLB cell types (Neng et al., 2013; W. Wang et al., 2015; Yi et al., 2023) or fragmented explant culture (Davis et al., 1995; Mali et al., 1997; J. Zhang et al., 2021). While these methods provide a platform for investigation, they do not examine the SV as a whole tissue. This could lead to an oversight of key information that could contribute to our understanding of the SV and the evaluation of therapies. Recently, single cell RNA sequencing has allowed for cell-specific characterization of the whole SV in both normal and hearing loss mouse models (Gu et al., 2020; Jean et al., 2023; Korrapati et al., 2019; Milon et al., 2021; Taukulis et al., 2021). However, the molecular underpinnings of SV development, maintenance and degeneration have yet to be fully elucidated.

We sought to provide a deeper understanding of the SV by contributing to the advancement of *in vitro* and bioinformatic studies. Whole organ explant cultures provide the advantage of preserving architectural and molecular integrity, which can better inform understanding of the organ *in vivo* and allow for more comprehensive *in vitro* studies including damage paradigms and therapeutic interventions. Therefore, we developed an organotypic explant model of the SV for both neonatal and adult mice. We showed that our *in vitro* system recapitulated SV proliferation *in vivo*, and conducted pharmacological studies using our system, demonstrating a robust, representative, and reproducible model to study the SV. To then understand the transcriptomic landscape of the SV further, we used single-cell RNA sequencing and compared the neonatal and adult SV. We revealed significant differences in gene expression patterns and bioinformatically characterized the proliferative properties of SV cell types. We further identified key genes, transcription factors, pathways, and cell-to-cell interactions unique to each stage that may play a role in SV proliferation, development, and maintenance. Overall, our novel experimental platform and single-cell RNA sequencing data provide new knowledge and insights into understanding the SV in the pursuit of developing biological solutions for SV-associated hearing loss.

## MATERIALS AND METHODS

### Animals

Care and euthanasia of male and female CD-1 mice (Charles River Laboratory) used in this study was approved by the Institutional Animal Care and Use Committee regulations at Sunnybrook Research Institute in Toronto, Canada. CBA/J mice were purchased from JAX (Stock No. 000656) and postnatal day 1 (P1) and P30 mice were used for single cell RNA sequencing experiments. The animal study was reviewed and approved by Animal Care and Use Committee of the National Institute of Neurological Diseases and Stroke and the National Institute on Deafness and Other Communication Disorders, National Institutes of Health.

### Stria Vascularis Organotypic Explant Cultures

Inner ears were dissected from neonatal and adult mice in ice cold HBSS with 1 % HEPES. In neonatal pups, the cartilaginous membrane over the cochlea was carefully removed, and in adults, the bone was carefully chipped off, to expose the cochlear tissue. The cochlear duct was removed from the modiolus, and the lateral wall was removed from the sensory epithelium. Then, the SV was gently teased away from the spiral ligament. SV were cultured in 2.5 % FBS media in a 10 mm glass bottom dish coated with Matrigel for a total of 72 hours (h) at 37° C. All experiments were stopped after 72 h, then explants were rinsed with PBS and fixed with 4% paraformaldehyde.

### In Vitro Proliferation Assays

The thymidine analog 5-bromo-2’-deoxyuridine (BrdU; BD Biosciences 550891) was used to assess cell proliferation *in vitro*. SV from postnatal day (P) 0-1 mice were exposed to 3.5 µg/mL BrdU for the following durations: 1.5 h (n = 11), 5 h (n = 10), 9 h (n = 10), 24 h (n = 13), 48 h (n = 12), and 72 h (n = 14). Proliferation was also assessed throughout postnatal age, at P0-1 (n = 10), P7-8 (n = 12), and P30-35 (adult; n = 6).

### Wnt Inhibition

SV were cultured in the presence of the β-Catenin/Tcf Inhibitor, FH535, at the following concentrations: 1 µM (n = 9), 2.5 µM (n = 10), 5 µM (n = 10), or 10 µM (n = 9); EMD Millipore 219330). DMSO was administered as the vehicle control (n = 9). All explants were cultured with 3.5 µg/mL BrdU for 72 h.

### Real-time quantitative PCR (RT-qPCR)

RT-qPCR experiments were performed to compare *mKi67* gene expression between neonatal and adult SV. Eight SV were pooled per sample, and experiments were performed using three biological replicates. RNA was extracted using the RNAqueous™-Micro Total RNA Isolation Kit (Invitrogen™ AM1931), and cDNA was transcribed using the High-Capacity RNA-to-cDNA™ Kit (Applied Biosystems™ 4387406) according to manufacturer’s instructions. RT-qPCR was performed using the TaqMan™ Fast Advanced Master Mix (Applied Biosystems™ 4444556) and run on the Applied Biosystems™ QuantStudio™ 5. mKi67 was tested in triplicate. The TaqMan probes used for RT-qPCR gene expression assays were GAPDH (Mm99999915_g1) and mKi67 (Mm01278617_m1).

### Tissue Cryosection

Inner ears were dissected from P0-1 or P30-35 mice and immediately fixed in 4% paraformaldehyde overnight at 4° C. Adult mouse temporal bones were decalcified in Osteosoft® (Millipore Sigma 1017281000) for 24-48 h at 37° C. Temporal bones were cryoprotected in 10%, 20% and 30% sucrose steps before being embedded in Tissue Tek® O.C.T. Compound (Sakura 4583). Tissues were sectioned at 10 µm thickness on Superfrost™ Plus Microscope Slides (Fisher Scientific 12-550-15).

### Immunofluorescence

Cryosections were permeabilized with 0.5% Triton X in PBS and quenched with 0.3 M Glycine in 0.5% Triton X in PBS. Antigen retrieval was performed prior to staining for BrdU using 1 N HCl for 30 min at room temperature. Sections were then blocked in 10% Donkey Serum for 1 h at room temperature. The following primary antibodies were used: Purified mouse anti-BrdU (1:250; BD Biosciences 555627), rabbit anti Ki67 (1:500, Abcam ab15580), rabbit anti Cldn11 (1:250; Santa Cruz sc-25711), rabbit anti Kir4.1 (1:250; Alomone Labs APC-035), and guinea pig anti Kcnq1 (1:250; Alomone Labs APC-022-GP). Griffonia (Bandeiraea) Simplicifolia Lectin I (GSL I, BSL I), Fluorescein (1:250; Vector Laboratories FL-1101-2), or Rhodamine (Vector Laboratories RL-1102-2) was used to stain blood vessels. DAPI was used as a counterstain for nuclei (1:1000, Sigma-Aldrich D9542).

### Microscopy

Images were acquired using a Nikon A1R Laser Scanning Confocal Microscope or a Nikon Eclipse Ti. Z-projections covering the depth of the explant or section were captured and converted into a maximum intensity projection. Large image acquisition was also performed with optimal path stitching and 15% overlap. For explant experiments, signal gain was determined using the control sample and then applied to the treated sample and were kept consistent between experimental repeats.

### Cell Quantification

Quantification was performed using FIJI. In all experiments, proliferation was quantified by calculating the percentage of BrdU+ cells among all DAPI labelled nuclei across the entire explant.

### Stria Vascularis Sample Preparation for scRNA sequencing

SV tissue samples were prepared as previously described (Korrapati et al., 2019). Briefly, mice were sacrificed, and inner ears from a total of four P1 mice and four P30 mice were collected. The SV was micro-dissected from the spiral ligament and lysed in either 0.5 mg/ml trypsin at 37°C for 7 min for P1 tissue, or 400-600 units/mL accutase at 37°C for 25 min for P30 tissue. Media was gently replaced with 5% FBS in DMEM to stop lysis, and the tissue was triturated and then filtered using a 20 µm filter (pluriSelect Life Science, El Cajon, CA, United States), and the cells were kept on ice for 35 min. The cell pellet was then suspended in 50 µL of the filtered media and cell counts were performed using a Luna automated cell counter (Logos Biosystems, Annandale, VA, United States). A cell density of 1 x 10^6^ cells/ml was used to load onto the 10X Genomics chip.

### 10x Chromium Genomics Platform

Single cell captures were performed following manufacturer’s recommendations on a 10x Genomics Controller device (Pleasanton, CA, United States). The number of captured cells per sample were as follows: 2259 cells, 111,122 mean reads per cell, 2422 median genes per cell from sample P1_s353n, 7790 cells, 16,260 mean reads per cell, 1859 median genes per cell from sample P1_s405n, and 5206 cells, 26,580 mean reads per cell, 1042 median genes per cell from sample P30_accu. Library preparation was performed according to the instructions in the 10x Genomics Chromium Single Cell 3’ Chip Kit V2. Libraries were sequenced on a HiSeq 1500 or Nextseq 500 instrument (Illumina, San Diego, CA, United States) and reads were subsequently processed using 10x Genomics CellRanger analytical pipeline using default settings and 10x Genomics downloadable mm10 genome. Dataset aggregation was performed using the cellranger aggr function normalizing for total number of confidently mapped reads across libraries. After filtering steps, we had 1973 cells for sample P1_s353n, 5724 cells for sample P1_s405n, and 2118 cells for sample P30_accu.

### Single Cell RNA Seq Data Preprocessing

Quality control was conducted using the following parameters: nFeature_RNA >900 & nFeature_RNA <5000 & percent.mt <15. Doublet detection was conducted with scDblFinder using the default parameters without clustering information. The raw gene expression matrix was normalized using the scTransform function in Seurat (Choudhary & Satija, 2022), and the P1 and P30 datasets were integrated after batch correction using the reciprocal principal component analysis (RPCA) method.

### Dimensionality reduction, Clustering and Data Visualization

Dimensionality reduction was performed via UMAP using the first 20 principal components. Clustering was performed in smart local moving algorithm in Seurat (Waltman & Van Eck, 2013). Clusters were annotated using known published markers for each cell type (Jean et al., 2023).

### Differential Expression Analysis and Functional Enrichment Analysis

Differential gene expression was performed in Seurat using the Wilcoxon Rank Sum test. Average log2fc > |1|, adjusted P-value < .05. Functional enrichment analysis was performed using gProfiler v. e109_eg56_p17_1d3191d. Genes that were upregulated in P1 and P30 were run as separate queries per cell type, filtering out ribosomal genes. Significance threshold was set to .05 and multiple tests were corrected using the Bonferroni correction method.

### CellChat Analysis

We performed intercellular communication (ligand-receptor) analysis for P1 and P30 datasets, respectively, using CellChat (v1.5.0, default parameters; (Jin et al., 2021)). Trimean algorithm was used to infer the communication network, including signalling pathway and ligand-receptor pairs information. We visualized sources and targets of each signal as circus plots via pycirclize (https://github.com/moshi4/pyCirclize)

### Trajectory Analysis

We applied trajectory analysis for the fibrocyte and basal cell clusters using Slingshot v1.6.0 (Street et al., 2018) and set the fibrocyte cluster as the starting point. We visualized the expression changes (normalized values) of marker genes along with the trajectory.

### Experimental Design and Analysis

Statistical analyses were performed using GraphPad-Prism software. For *in vitro* proliferation assays and the Wnt inhibition study, a One-way ANOVA with Tukey’s Multiple Comparisons Test was performed for all experiments with the significance threshold set to p < 0.05. For RT-qPCR experiments, a two-tailed Student’s t-test was performed for each gene between ages or conditions with the significance threshold set to p < 0.05. All data are represented as mean and standard error of the mean (SEM). All n-values presented refer to number of SV. All p-values are presented in figure legends.

### Data Availability

The single cell sequencing data generated in these studies have been deposited in the Gene Expression Omnibus (GEO) database (GEO Series accession ID: GSE262019, https://www.ncbi.nlm.nih.gov/geo/query/acc.cgi?acc=GSE262019)

## RESULTS

### Establishing an *in vitro* culture model for the SV

The development of a whole-tissue culture model will enable comprehensive studies for the SV and create a platform for high throughput testing of therapeutics. With these goals in mind, we established an organotypic explant culture model using neonatal and adult mice to study the whole SV. In brief, SV and associated vasculature were dissected from base to apex from P0-P35 CD1 mice and cultured on Matrigel-coated plates. All stages were cultured for 72 h before fixation with 4% PFA. We initially observed that the P0-1 SV had growth and spread of cells while the P30-35 SV did not (Fig. 1C). We hypothesized that this could be a result of cell proliferation, and to assess this, we cultured P0-1 SV for 72 h and exposed the explants to 3.5 μg/mL BrdU for different durations within the culture period (Fig 2A-C’). We detected BrdU^+^ nuclei within 1.5 h in culture and significant accumulation of BrdU within 24 h (F(5, 64) = 56.23, p < 0.0001; Fig. 2D). Together, our results demonstrated that the P0 SV exhibits active and persistent proliferation *in vitro*.

**Figure 2.**
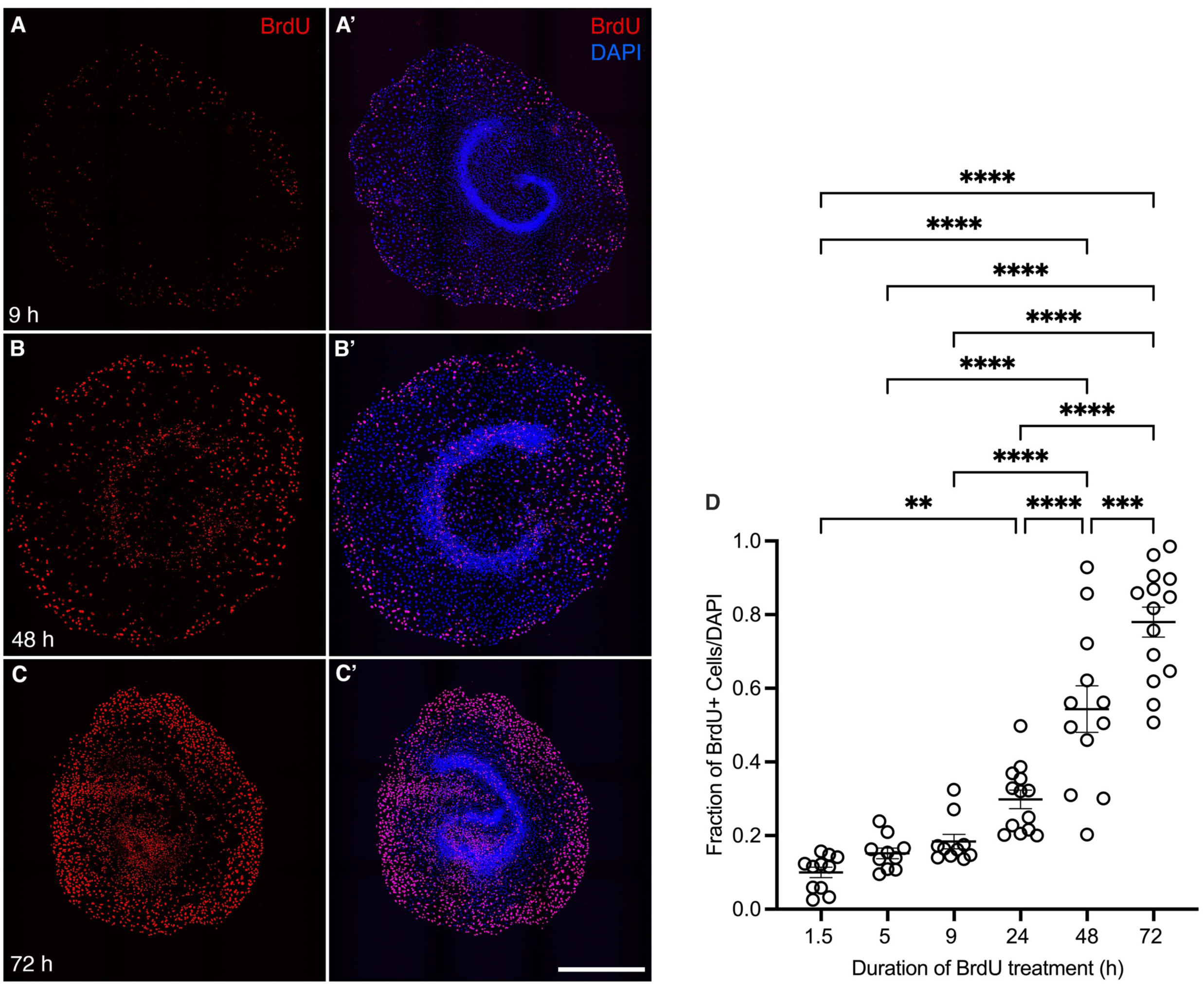
Proliferation of the P0 SV is ongoing throughout 72 h in culture. (A-C) Representative images of P0 SV proliferation across different timepoints over 72 h with the proliferation marker, BrdU (red). (A’-C’) Merged images of BrdU^+^ nuclei and DAPI stained nuclei (blue). Scale bar: 1mm. (D) Proliferation quantified as the fraction of BrdU^+^ nuclei over total DAPI stained nuclei. A one-way ANOVA was performed, F(5, 64) = 56.23, p < 0.0001. Tukey’s multiple comparisons test was performed for post-hoc analysis. **p = 0.0044, ***p = 0.001, ****p < 0.0001. Graph shows mean with SEM.

### The neonatal SV is highly proliferative *in vitro*, and proliferation decreases during postnatal development

We next determined the proliferative capacity of the SV during postnatal development. We cultured SV from P0-1, P7-8, and P30-35 mice and observed a significant decrease in proliferation as measured by the proportion of BrdU^+^ cells in relation to DAPI-labeled cells at P7-8 (p = 0.02) and P30-35 (p < 0.0001) compared to P0-1 (Fig. 3A-D). This indicated that while the SV retains its proliferative capacity at neonatal stages, proliferation decreases with age.

**Figure 3.**
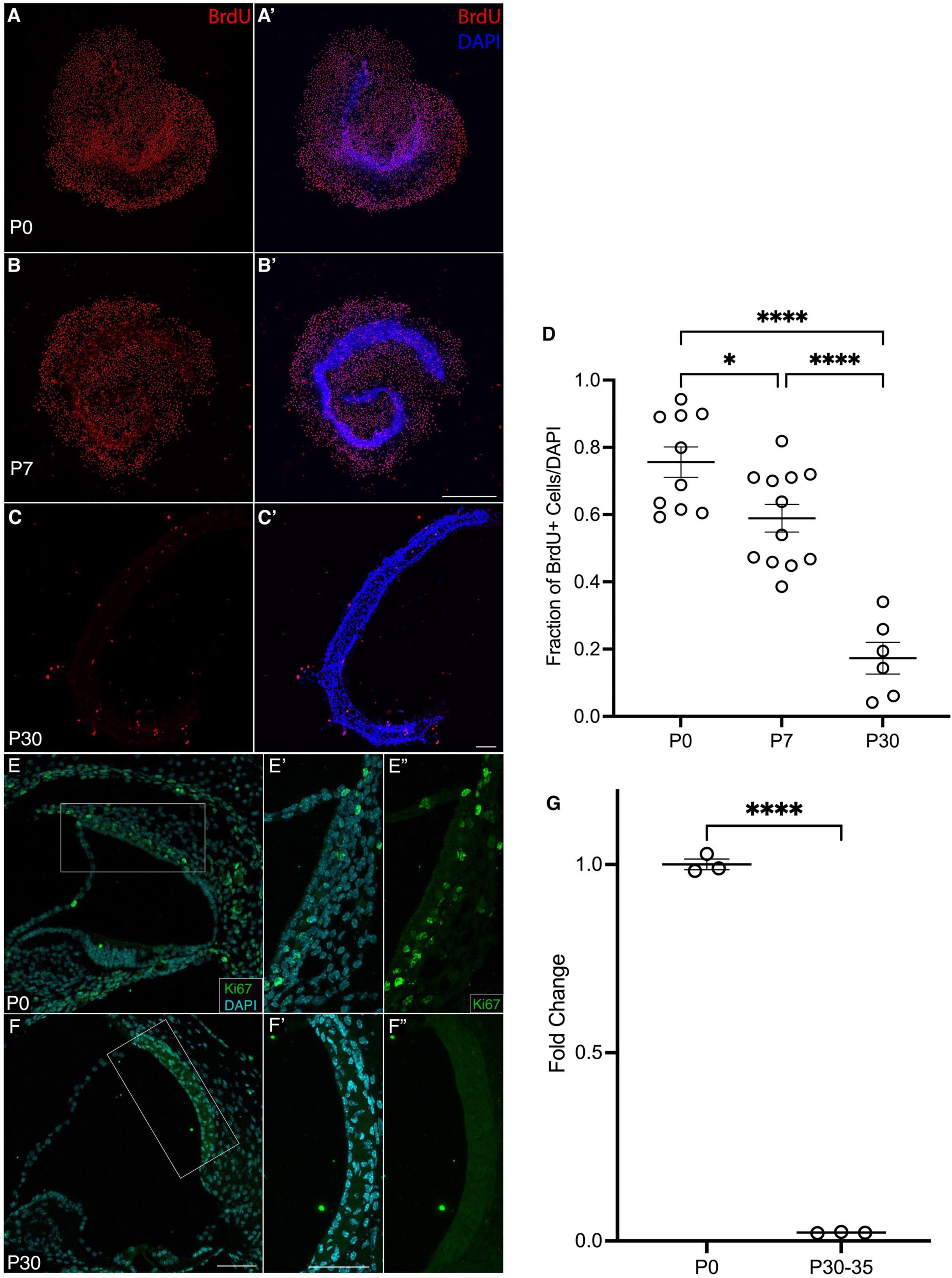
Proliferation of the stria vascularis decreases with age. (A-C) Representative images of organotypic stria vascularis explants at P0, P7 and P30 cultured with the proliferation marker, BrdU (red), over 72 h. (A’-C’) Merged images of BrdU^+^ nuclei and DAPI stained nuclei (blue). Scale bars: 1 mm in A-B’; 200 µm in C-C’. (D) Quantification of BrdU^+^ nuclei. A one-way ANOVA was performed, F(2, 25) = 34.15, p < 0.0001. Tukey’s multiple comparisons test was performed for post-hoc analysis. (E-F”) Cryosections of the basal turn of the P0 and P30 cochlea were stained with Ki67 (green), Lectin (magenta) and DAPI. Scale bars: 100 µm for low magnification images; 50 µm in high magnification images. (G) Quantification of Ki67 in P0 and P30 SV using RT-qPCR. A two-tailed unpaired t-test was performed, t(4) = 70.16, p < 0.0001. *p = .0235, ****p < .0001. All graphs show mean with SEM.

Next, we examined whether the proliferative effect we observed *in vitro* recapitulated the behavior of the SV in the ear. We validated proliferation *in vivo* by immunolabelling for Ki67, an endogenous marker of proliferation (Miller et al., 2018; Sobecki et al., 2016) in cryosections of P0 and P30 mouse cochleae. In agreement with published reports (Renauld et al., 2022), we observed Ki67^+^ cells at P0 in the intermediate cell layer near the vasculature but not at P30 (Fig 3E-F”). Furthermore, using RT-qPCR, we quantified a significant decrease in *Ki67* mRNA expression in P30-35 SV compared to P0-1 SV (p < 0.0001; Fig3G). Taken together, the proliferative behaviour of the SV in our *in vitro* was comparable to *in vivo*, validating our system as a representative experimental platform to investigate the SV.

### Wnt/β-catenin signalling plays a role in SV proliferation

We then tested the utility of our *in vitro* system to pharmacological intervention by asking what molecular pathways were driving proliferation in the neonatal SV. A well-known pathway involved in proliferation and angiogenesis is the canonical Wnt/β-catenin (Wnt) signalling pathway (reviewed in Steinhart & Angers, 2018). We have previously shown that the canonical Wnt signalling pathway is highly active in the developing cochlea, and that activating Wnt signalling at embryonic and postnatal stages promotes proliferation of cochlear supporting cells (Jacques et al., 2013; Samarajeewa et al., 2018). We have also reported that Wnt signalling components are expressed in the SV (Geng et al., 2016; Korrapati et al., 2019; Samarajeewa et al., 2019). To examine the role of Wnt signaling in SV proliferation, we used a pharmacological inhibitor of Wnt signalling called FH535 which targets the TCF/LEF transcription factors responsible for regulating downstream Wnt target genes. We administered FH535 at 1, 2.5, 5, and 10 µM for 72 h on P0-1 cultured SV and found that proliferation significantly decreased in cultures treated with 5 µM (p = .0003) and 10 µM (p < 0.0001) FH535 compared to DMSO controls (Fig. 4A-D). These results showed that our culture model is robust to experimentation including pharmacological intervention and indicated that canonical Wnt signalling is at least in part responsible for regulating proliferation in the SV.

**Figure 4.**
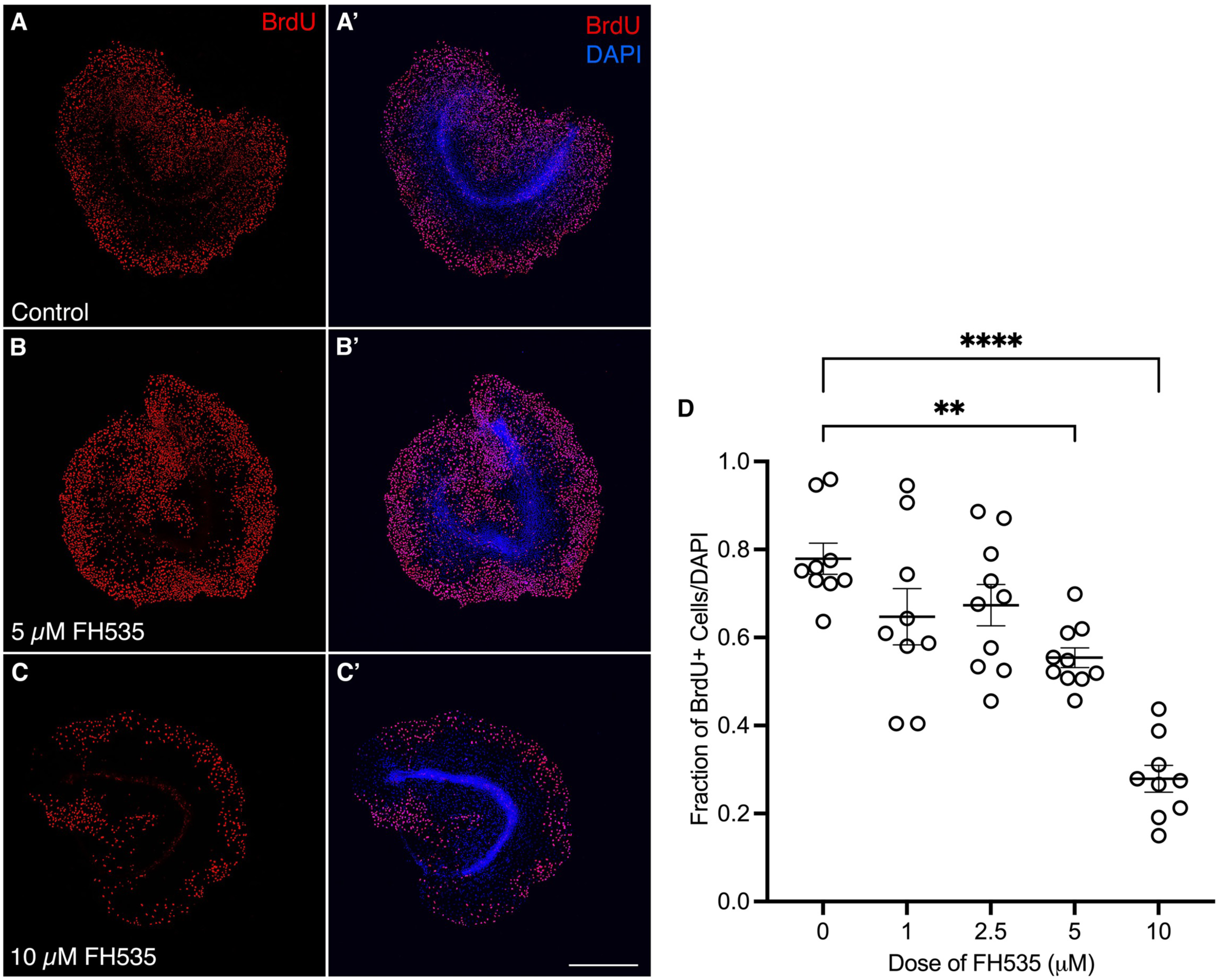
Inhibition of Wnt/β-catenin signaling significantly reduces proliferation of the neonatal stria vascularis. (A-C) Representative images of organotypic P0-P1 stria vascularis explants cultured with either DMSO (control) or FH535, in the presence of BrdU (red) over 72 h. (A’-C’) Merged images of BrdU^+^ nuclei amongst DAPI stained nuclei (blue) in control and FH535 treated cultures. Scale bar: 1 mm. (D) Quantification of BrdU^+^ nuclei amongst DAPI. A one-way ANOVA was performed, F(4, 42) = 19.74, p < 0.0001. Tukey’s multiple comparisons test was performed for post-hoc analysis. **p = 0.004, ****p < .0001. Graph shows mean with SEM.

### Profiling the P1 and P30 transcriptome using single cell RNA sequencing

To gain a further understanding of the molecular differences between the neonatal and adult SV and their neighbouring cell populations, we utilized single cell RNA sequencing. We collected samples from *in vivo* P1 and P30 CBA/J mice and processed them as previously described (Korrapati et al., 2019). We performed unsupervised bioinformatic clustering using Seurat and resolved cluster identity using published cell type-specific markers (Jean et al., 2023; Korrapati et al., 2019). We then revealed defined marginal, intermediate, and basal cell clusters in both neonatal and adult SV and validated the expression of cell-type specific markers in these populations using immunofluorescence (Fig. 5A-B).

**Figure 5.**
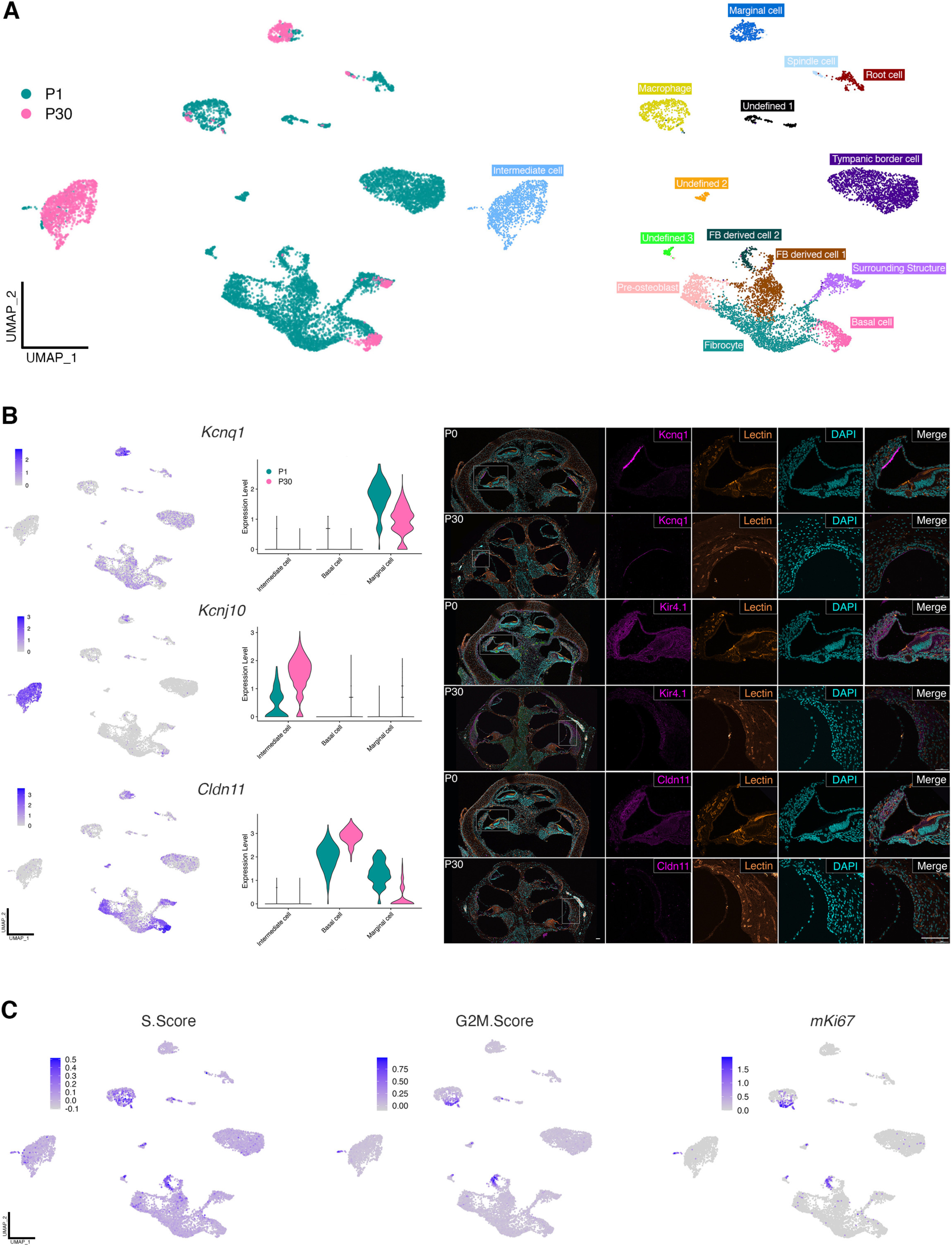
Annotation of cell types and proliferative clusters in P1 and P30 single cell RNA sequencing data. A) Integrated UMAP of P1 and P30 single cell RNA sequenced SV showing P1 and P30 sample distribution (A) among 15 cell clusters (B). C) UMAPS and violin plots of candidate marker genes for marginal, intermediate, and basal cells. D) Immunohistochemical validation of candidate marker genes in cryosections of P0 and P30 cochleae. Scale bars: 100 μm for low magnification images; 50 μm for high magnification images. E) UMAPs showing clusters in which cells undergo proliferation.

To annotate the remaining clusters in our dataset, we integrated P8, P12, and P20 single cell sequencing data from Jean et al., 2023. We identified root cells (solute carrier family 26 member 4 (*Slc26a4*) and epiphycan,( *Epyc*)), spindle cells (*Slc26a4* and annexin A1 (*Anxa1*)), macrophages (adhesion G protein-coupled receptor E1 (*Adgre1*) and macrosialin (*Cd68*)), fibrocytes (otospiralin (*Otos*)), tympanic border cells (elastin microfibril interfacer 2 (*Emilin2*) and palmitoleoyl-protein carboxylesterase (*Notum*)), pre-osteoblasts (RUNX family transcription factor 2 (*Runx2*) and distal-less homeobox 5 (*Dlx5*)), two mixed populations with endothelial and pericyte characteristics (which we labelled fibrocyte (FB) derived cell 1 and 2; Fms related receptor tyrosine kinase 1 (*Flt1*), endothelial cell adhesion molecule (*Esam*), Platelet-derived growth factor receptor β (*Pdgfrb*), and regulator of G protein signaling 5 (*Rgs5*)) and cells from the surrounding structure (odd-skipped related transcription factor 2 protein (*Osr2*) and chordin-like protein 1 (*Chrdl1*)). We also had three smaller clusters with unknown identity, which we have marked as “unidentified” (Fig. 5A).

To unveil proliferative differences between the P1 and P30 SV, we identified proliferating cell populations using S.Score and G2M.Score with Seurat, which calculates cell cycle scores using known markers of S phase and G2/M phase of the cell cycle. We identified that of the three main cell types, a subset of intermediate cells undergo proliferation at P1, which corresponds with the expression of the proliferation marker, *Ki67* (Fig. 5C).

We then sought to gain a deeper understanding of the specific cell types of the SV and surrounding cells. As fibrocytes are closely associated with basal cells, we were interested in understanding their relationship. Fibrocytes regulate ion homeostasis alongside the SV and play a major role in cochlear blood flow regulation, immune response, and recovery from trauma (reviewed in Furness, 2019; Peeleman et al., 2020). Fibrocytes proliferate after injury even in adult animals (Kamiya et al., 2007; Stevens et al., 2014), which makes them a very interesting candidate population to study lateral wall regeneration. We performed a pseudotime analysis and identified a clear trajectory of cell development from the fibrocytes to the basal cells from P1 to P30 (Fig. 6A). We analyzed the top 300 genes along the trajectory to further understand the transitional process of basal cell development. In accordance with previous reports (Trowe et al., 2011), we identified some genes that are involved in mesenchymal-to-epithelial transition of the fibrocytes into basal cells: expression of *Vim* (vimentin), a mesenchymal marker, decreased along pseudotime, while the expression of *Cdh1* (E-cadherin), an epithelial marker, increased. We also identified that expression of the gap junction protein genes, *Gjb2* (connexin 26) and *Gjb6* (connexin 30), increased in basal cells compared to fibrocytes, and expression of several genes from the collagen families decreased (Fig. 6A). Mutations of *Gjb2* and *Gjb6* are associated with the most common autosomal recessive hereditary hearing loss and deficiencies in one or both genes can cause significant reduction of the endocochlear potential (Chen et al., 2022; Mei et al., 2017). Collagen plays an important role in the spiral ligament (SL) as it helps form the extracellular matrix (Tsuprun & Santi, 1999) and thereby maintains the structural integrity of the lateral wall. Thus, our psuedotime analysis provides a molecular landscape of the distribution of functional genes that both distinguish and conjoin the SV and the SL.

**Figure 6.**
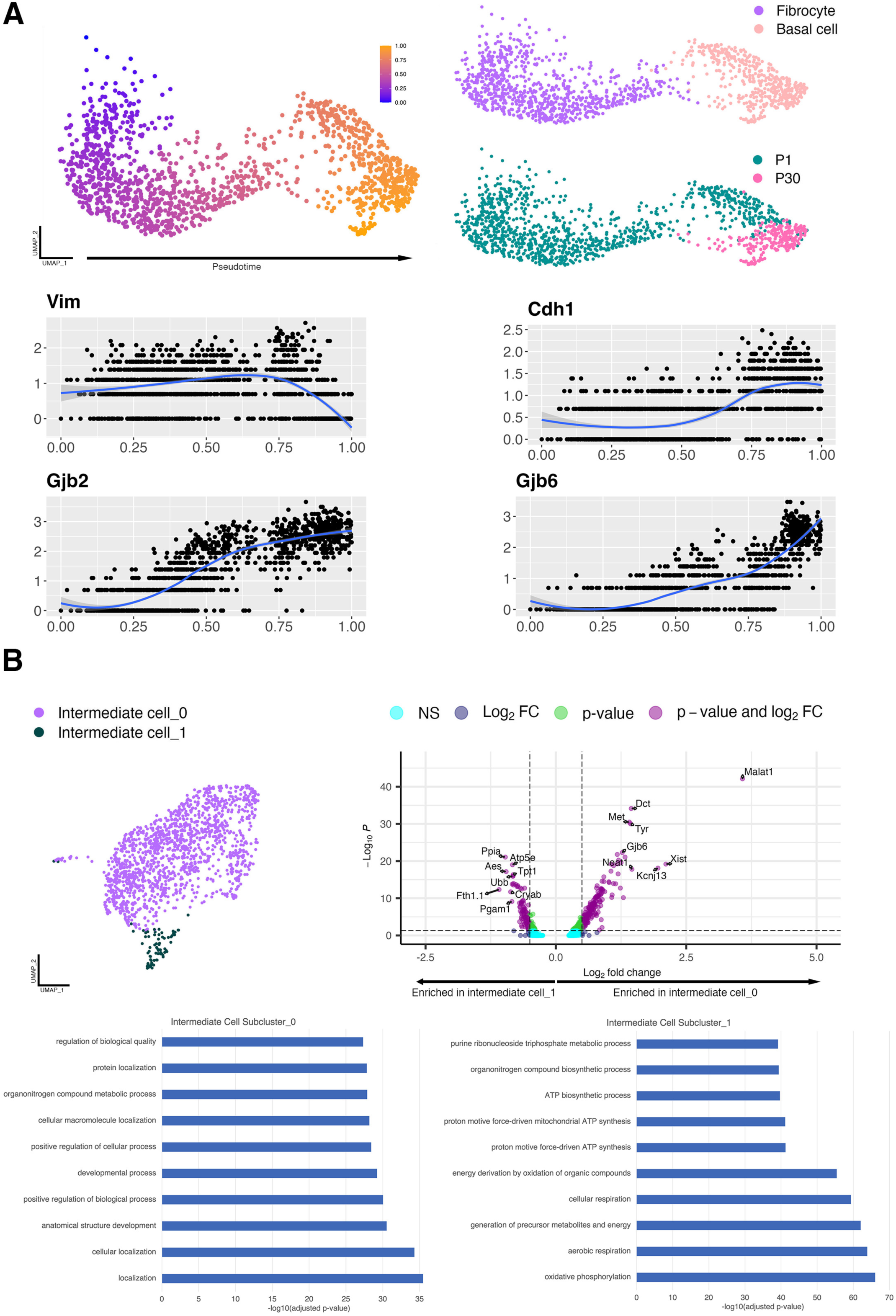
Subclustering and trajectory analyses on intermediate and basal cells. A) Pseudotime analysis showing fibrocyte to basal cell trajectory, and UMAPs displaying fibrocyte and basal cell subclusters and sample stage distribution. Expression of different genes in fibrocytes and basal cells along pseudotime. B) UMAP displaying intermediate cell subclusters and volcano plot showing differential gene expression between subclusters.

We further characterized the intermediate cells of the SV, as their exact identity is unclear. Intermediate cells are melanocytes that arise either directly from the neural crest, or indirectly from Schwann cell precursors, which are a population of neural crest cells which emigrate to the peripheral nerve early in development (Adameyko et al., 2009; Aoki et al., 2009; Bonnamour et al., 2020). A majority of intermediate cells are derived from Schwann cell precursors (Bonnamour et al., 2020; Renauld et al., 2022). We subclustered the intermediate cell population to determine if we could identify these two populations. Here, we identified two subclusters of intermediate cells, which we labelled subcluster_0 and subcluster_1 (Fig. 6B). Interestingly, our data showed that marker genes for subcluster_0 include many, if not all, of the recognized canonical markers of intermediate cells, including *Kcnj10* (potassium inwardly rectifying channel subfamily J member 10)*, Met* (met proto-oncogene)*, Dct* (dopachrome tautomerase), and *Plp1* (proteolipid protein 1). Genes upregulated in subcluster_1 include *Vegfb* (vascular endothelial growth factor B)*, Scn1b* (sodium voltage-gated channel beta subunit 1)*, Mt1* (metallothionein 1), and *Mapk3* (mitogen activated protein kinase 3). GO biological process terms associated with subcluster_0 included cellular localization and anatomical structure development, whereas subcluster_1 included oxidative phosphorylation and generation of precursor metabolites and energy. Although more investigation is required, these results may begin to elucidate the molecular identity of the intermediate cell subtypes and their origins.

### Differential gene expression analysis reveals transcriptional differences between the neonatal and adult SV

To examine the transcriptional differences between the neonatal and adult SV, we performed differential gene expression analysis comparing the P1 to P30 SV for marginal, intermediate, and basal cells (Fig. 7A-C). We were particularly interested in genes that were uniquely upregulated at either P1 or P30, or in one specific cell type. Among the top significantly enriched genes in P1 marginal cells, the ion channels *Dpp10* (dipeptidyl peptidase 10) and *Cacnb2* (calcium voltage-gated channel auxiliary subunit beta 2) were unique to this cluster. *Dpp10* is a potassium channel ancillary subunit and is known to associate with the voltage-gated potassium channel family, Kv4 (Li et al., 2006) and *Cacnb2* has been previously reported in the inner hair cells and is required for normal development and hearing function (Neef et al., 2009). Similarly, in P1 intermediate cells, the ion channel *Trpm1* (transient receptor potential cation channel subfamily M member 1) and the melanocyte-specific transmembrane protein *Pmel* (premelanosome protein) were upregulated. We further examined transcription factors that were differentially expressed between P1 and P30 in each cell type. There were fewer transcription factors expressed at P30 than at P1 in marginal and intermediate cells, but not basal cells. We again observed genes that were uniquely upregulated. For example, *Dach1* and *Meis1* are two transcription factors only enriched in P1 marginal cells, and they have been previously known to be involved in regulatory processes of the SV (Miwa et al., 2019; Nabec et al., 2022). Table 1 provides a complete list of all significant uniquely expressed genes, including transcription factors and cofactors, expressed at each age and cell type.

**Figure 7.**
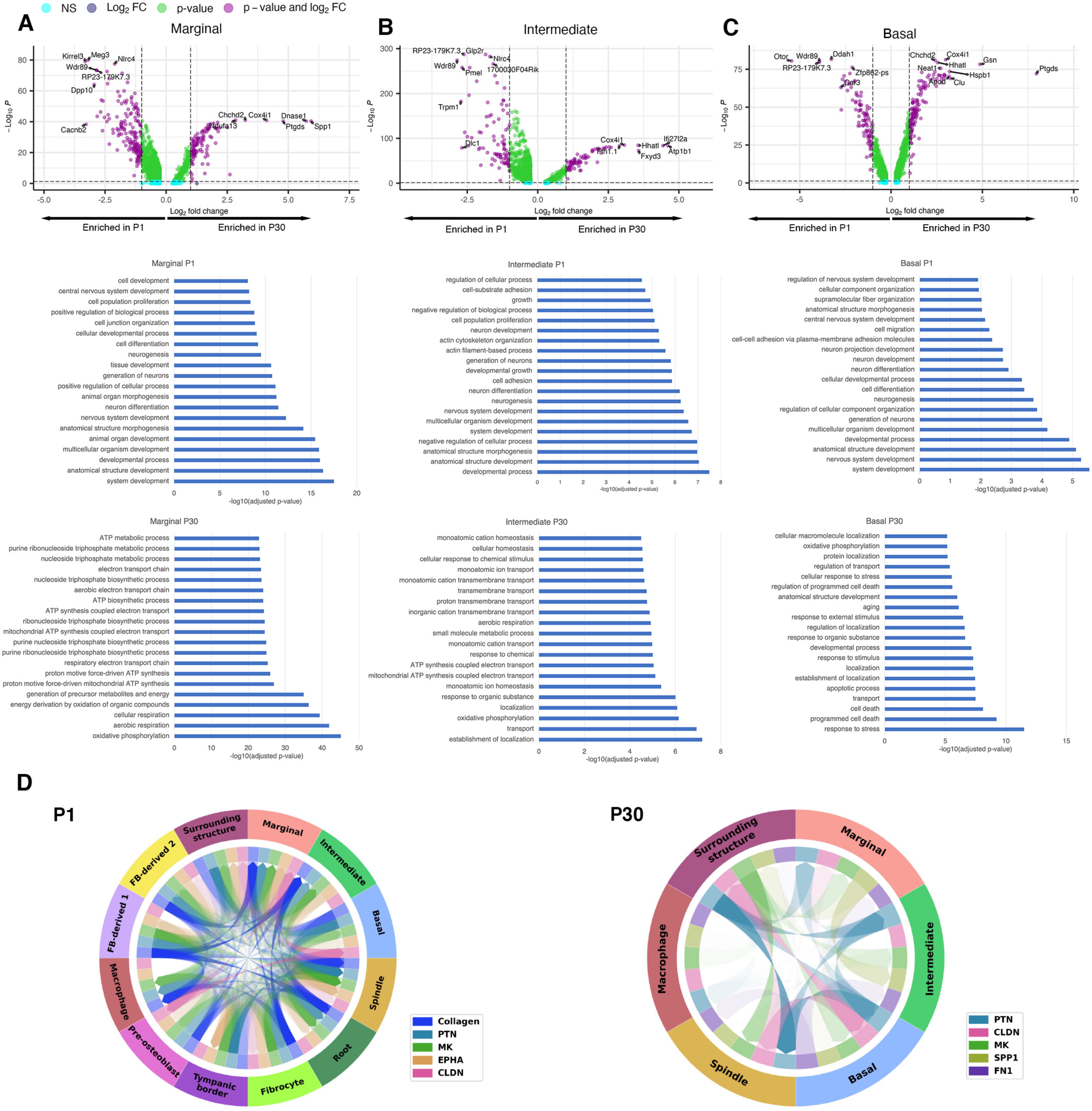
Differential expression and CellChat analyses reveal molecular differences between P1 and P30 SV. Differential gene expression analysis (volcano plots on the left) and GO biological processes (bar plots on the right) determined from P1 and P30 SV in A) marginal cells, B) intermediate cells, and C) basal cells. D) Circos plots from CellChat analysis displaying top five pathways and interaction patterns in P1 and P30 cell types.

**Table 1.**
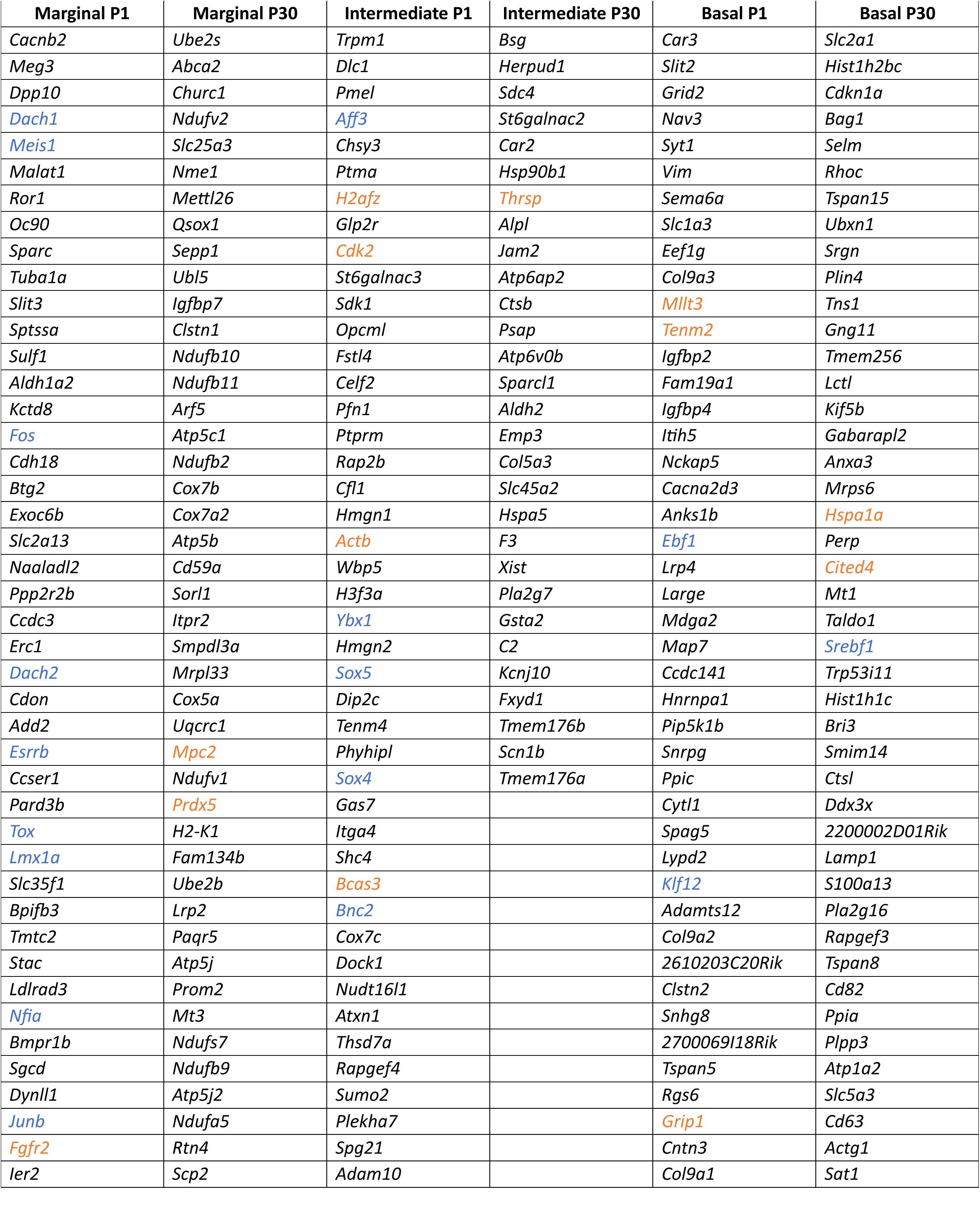
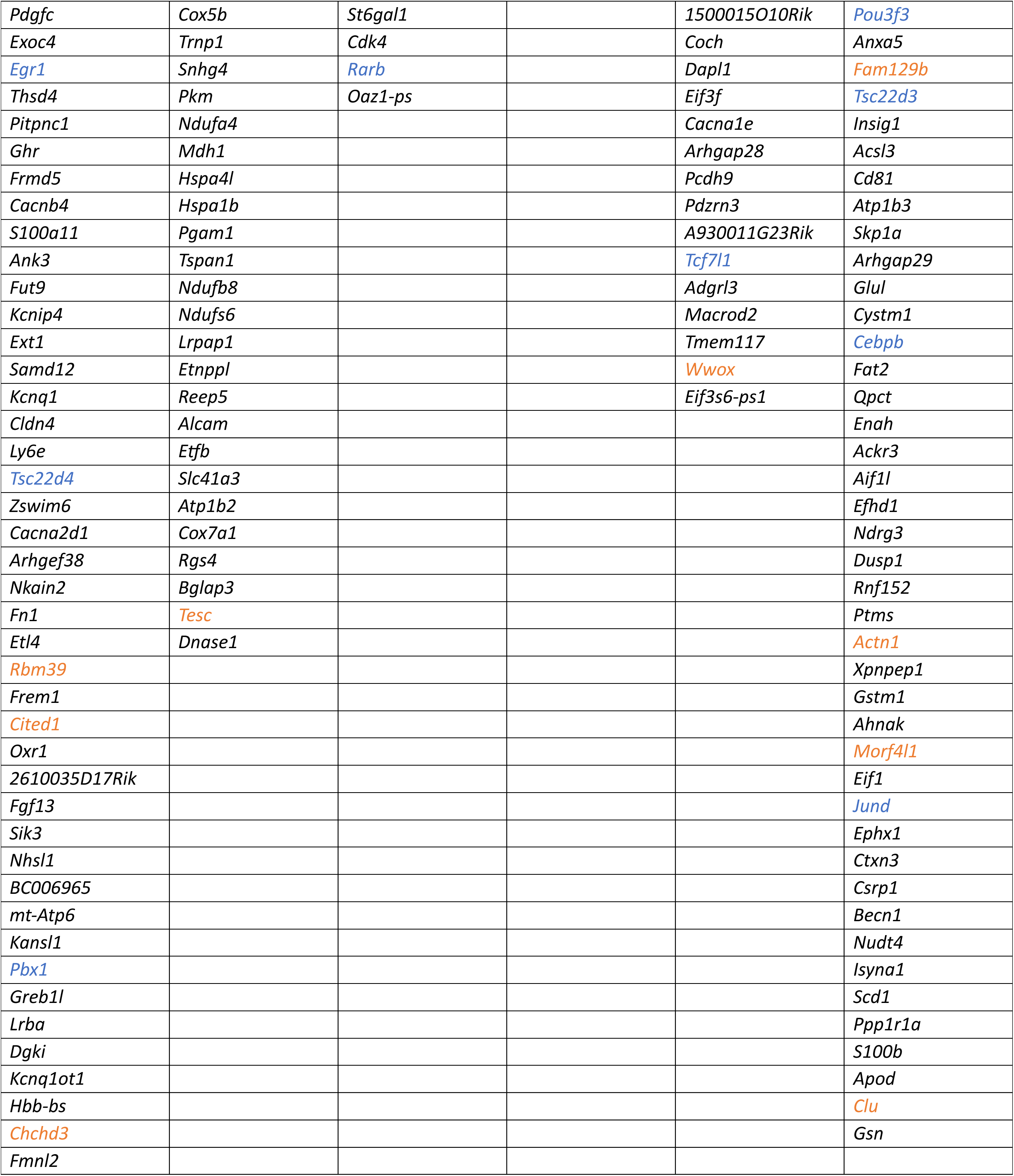
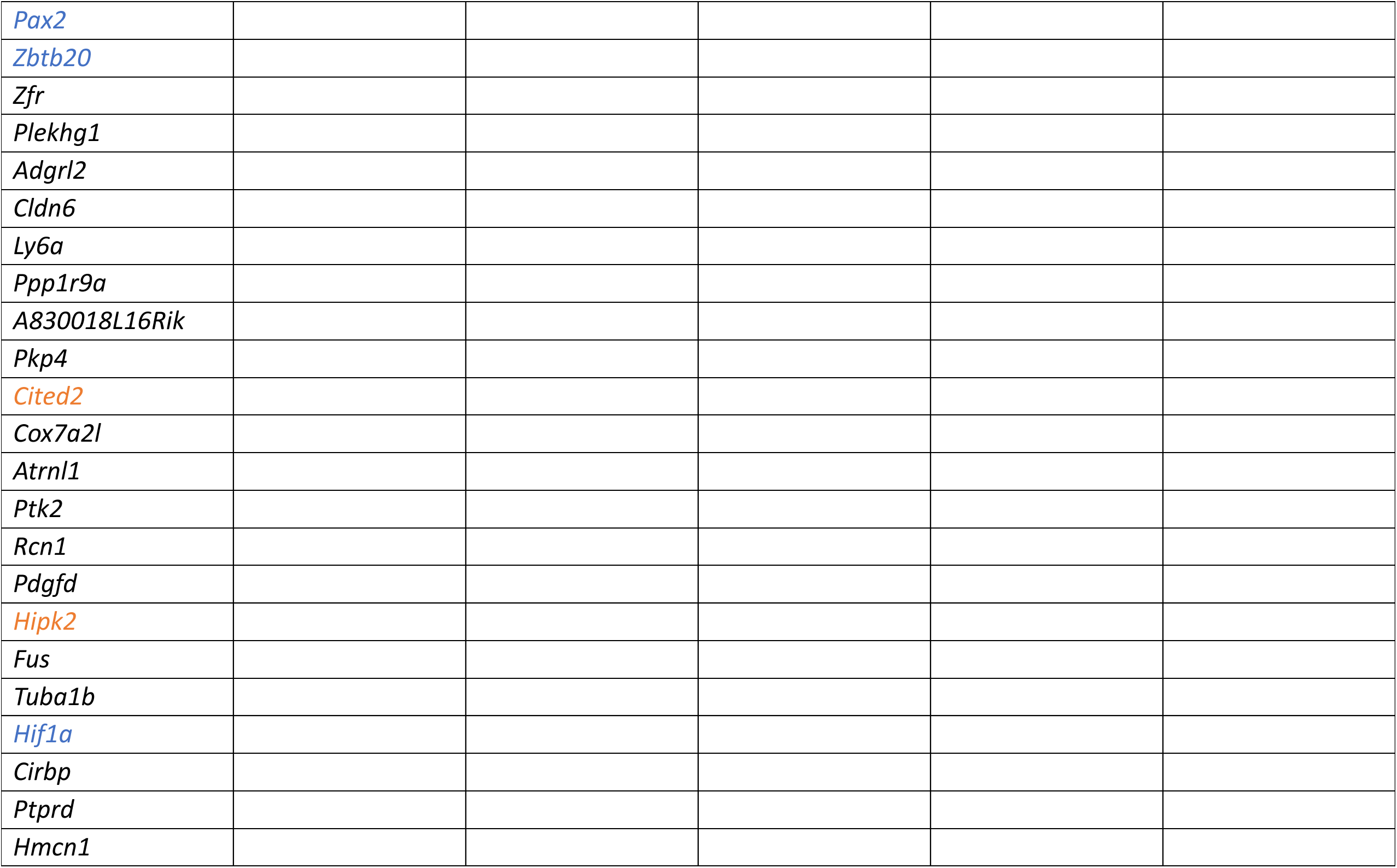
Unique differentially expressed genes in P30 vs. P1 SV cell types. Significant genes were determined using log2Fc > |1| and Padj < 0.05. Transcription factors (blue) and co-factors (orange) extracted using animalTFDB.

We then performed functional enrichment analysis using gProfiler, using all significantly upregulated genes at P1 and P30 in marginal, intermediate, and basal cells. We ran independent analyses for each cell type at each stage and observed that the top 20 enriched GO terms for biological processes in P1 were widely associated with development, while at P30 they were associated with SV function and maintenance (Fig. 7A-C). Terms associated with proliferation and differentiation were identified at P1: cell population proliferation (GO:0008283) in marginal and intermediate cells, cell differentiation (GO:0030154) in marginal and basal cells, neuron differentiation (GO:0030182) in intermediate cells, and cell migration (GO:0016477) in basal cells. Collectively, differential gene expression analysis coupled with functional enrichment analysis of single cell transcriptomes from both the proliferative neonatal and non-proliferative adult SVs enabled a more comprehensive examination of the changes underlying SV development.

### CellChat examines cell-cell communication and ligand-receptor interactions within the neonatal and adult SV

The previous analyses provided an understanding about cellular identity within the specific populations at neonatal and adult stages. To gain further insight into cell-cell interactions, we performed CellChat analysis. We revealed several pathways involved in forming the extracellular matrix, cell proliferation and differentiation, and maintaining tissue structural integrity (Fig. 7D). We observed that cell-cell interaction patterns and pathway signaling strength differed between the P1 and P30: in many cases, at P1, a signaling pathway was expressed in several different cell types, while at P30, expression becomes more specified to fewer cell types. We also identified some pathways that are specific to P1 and P30, for instance, Vcam signaling at P1, and Tweak signaling at P30.

At P1, the top five signaling pathways were Collagen, Ptn (pleiotropin), Mk (midkine), Epha, and Cldn (claudin), and at P30, they were Ptn, Cldn, Mk, Spp1 (secreted phosphoprotein-1), and Psap (prosaposin). Some proteins, such as collagen and claudin, have been studied in the SV. Collagens are important extracellular matrix proteins that maintain the structural integrity of the SV (Cosgrove et al., 1998, 2008; Dufek et al., 2020; Meyer Zum Gottesberge et al., 2008) and Cldn (Claudins) are cell adhesion proteins that maintain the SV by regulating tight junctions (Gow et al., 2004; Kitajiri et al., 2004). There is considerably less information regarding Ptn, Mk, Spp1, and Psap. Ptn is an evolutionarily conserved neurotrophic factor which shares homology with Mk and regulates developmental and angiogenic processes (reviewed in Papadimitriou et al., 2021). *Ptn* or *Mk* knockout in mice lead to a lack of Kir4.1 expression in intermediate cells (Sone et al., 2011). Epha is a receptor tyrosine kinase belonging to the ephrin receptor subfamily (Defourny, 2019). Spp1 is a signaling protein which plays a role in immune function in disease (reviewed in Yim et al., 2022). In the inner ear *Spp1* has been previously characterized as a type I vestibular hair cell marker and more recently identified in cisplatin-treated SV (McInturff et al., 2018; Taukulis et al., 2021). Finally, Psap is precursor protein to the sphingolipid activator protein family (saposins) and is a secreted neuro- and glio-protective protein (Meyer et al., 2014). Prosaposin has been identified in the organ of Corti of rats and mice (Akil et al., 2006), and in the basal cells of the rat SV (Terashita et al., 2007). *Psap* knockout mice show impaired hearing (Akil et al., 2006) but more investigation is required to understand its role in the SV. Cell-cell communication analysis of the SV at neonatal and adult stages has provided more information about the intricate interactions within the SV and has revealed pathways that may have been previously overlooked.

## DISCUSSION

The goal of this study was to one, provide an accessible platform to study the SV, and two, provide a comparative profile of the molecular differences between the neonatal and adult SV. We used whole-organ *in vitro* explants of neonatal and adult SV to show the proliferative differences throughout postnatal development, and we used single cell RNA sequencing to further compare the transcriptome profiles of neonatal vs. adult SV. Our comprehensive analyses provide in-depth insights into the molecular composition of the SV and reveal key genes and pathways that contribute to SV-associated hearing loss pathologies.

We are the first to culture whole SV in both neonatal and adult mice. One advantage of our protocol is that we preserve the cellular architecture of the SV to study the interactions between cell types along the basal-to-apical axis of the tissue. Our system can be used to investigate different biological pathways involved in SV development, regulation, and disease, compare age-related differences in the SV, as well as test the effectiveness of therapeutic candidates.

We investigated SV proliferation *in vitro* using the thymidine analog, BrdU, and *in vivo* using Ki67. Consistent with previous reports using neonatal fragmented SV cultures, we observed proliferative cells that migrated out from the explanted neonatal tissue (Mou et al., 1997) which decreases during postnatal development (Renauld et al., 2022). We identified that a molecular mechanism regulating SV proliferation was Wnt/β-catenin signaling which is a prominent pathway involved in proliferation (Díaz-Coránguez et al., 2020; Steinhart & Angers, 2018). Wnt/β-catenin signaling is initiated when Wnt ligands bind to Frizzled receptors and Lrp5/6 co-receptors. This triggers the events that lead to the accumulation and subsequent translocation of intracellular β-catenin to the nucleus. Nuclear β-catenin then binds to the TCF/LEF transcription factors to activate the transcription of downstream Wnt target genes. We showed that inhibiting TCF/LEF transcription factors results in a significant decrease in proliferation in neonatal cultures, indicating that the Wnt/β-catenin signaling pathway plays a role in the proliferation of the SV. This suggests that Wnt/β-catenin signaling may be a therapeutic target for regenerative SV therapies.

We identified differentially expressed genes and intercellular interactions between these cell types in the neonatal and adult SV that may play a role in proliferation, differentiation, and development. *Dach1* is a transcription factor that we found to be upregulated in P1 marginal cells. Embryonic shRNA-regulated knockdown of *Dach1* results in a loss of intermediate cells, decreased endocochlear potential, and decreased hearing function in adulthood (Miwa et al., 2019). These results suggest that *Dach1* may be an important regulator of SV development. It also suggests that the association between marginal and intermediate cells may be an important factor in SV development. In accordance with this, we observed the upregulation of *Spp1* (secreted phosphoprotein 1) in P1 marginal and intermediate cells. *Spp1* has been characterized as a vestibular Type I hair cell marker using single cell RNA sequencing (McInturff et al., 2018). In the SV, the protein encoded by *Spp1*, osteopontin, is highly localized in the marginal cells and secreted into the cochlear fluids (Davis et al., 1995; Lopez et al., 1995).

Interestingly, *Spp1* is downregulated in the marginal cells of adult mice that have been treated with cisplatin, an ototoxic chemotherapeutic (Taukulis et al., 2021). These observations in adults coupled with our data in neonates suggest that *Spp1* may be a candidate gene to investigate SV and endocochlear potential development and is a therapeutic target for cisplatin-induced degeneration of the SV and hearing loss.

Meniere’s disease is another disorder that effects the SV. Meniere’s disease is a rare inner ear disorder characterized by endolymphatic hydrops and resulting in vertigo, tinnitus, and hearing loss. The etiology of Meniere’s disease is unclear. We identified that a potential therapeutic target for Meniere’s disease is the TWEAK signaling pathway. TWEAK (tumor necrosis factor-like weak inducer of apoptosis) is a member of the tumor necrosis factor superfamily and is a cytokine that has multiple functions in the body, including in inflammation, bone remodeling, angiogenesis, cellular adhesion, proliferation, differentiation, and apoptosis (Du et al., 2015; Ratajczak et al., 2022; Winkles, 2008). We identified that the TWEAK pathway is upregulated in the P30 SV. A study examining bilateral Meniere’s disease identified that a majority of patients carried a single nucleotide variant rs4947296 which regulates the TWEAK/Fn14 signaling pathway (Frejo et al., 2017). TWEAK/Fn14 signaling in individuals carrying the variant lead to the activation of both the canonical and non-canonical Nf-κB pathways with downstream inflammatory effects. While TWEAK signaling was identified to be a candidate therapeutic target for individuals carrying the variant, it was unclear where the site of inflammation was in the ear (Frejo et al., 2017). Our CellChat data suggests that the TWEAK signal is sent from the intermediate cells and the receptor is located on the marginal cells and the spindle cells. More information is needed to elucidate the role TWEAK signaling plays in the SV, but pinpointing the pathways involved in Meniere’s disease and the ligand-receptor interactions between the cell types can facilitate the development of effective therapies.

Identification of key genes and pathways also provides more insight into the characterization of each SV cell type. We were most interested in the intermediate cells because of their multi-faceted characteristics and their dual embryonic origin. Intermediate cells are primarily defined as melanocytes that exhibit macrophage characteristics (Shi, 2010; W. Zhang et al., 2012). Melanocytes are reported to originate from either directly from the neural crest or from a subset of early delaminated neural crest cells called Schwann cell precursors (Adameyko et al., 2009; Aoki et al., 2009; Bonnamour et al., 2020). From the neural crest, or nerve-derived Schwann cell precursors, melanoblasts form and migrate to their desired location in the body and form region-specific melanocytes depending on signaling pathways in the environment, such as KiT/Kitl or Et3/Ednrb (Adameyko et al., 2009; Aoki et al., 2009; Kaucka et al., 2021; Renauld et al., 2021; Yoshimura et al., 2013). Intermediate cells of the SV derive from these two embryonic populations (Bonnamour et al., 2020; Renauld et al., 2022), yet the exact nature of the two intermediate cell populations remains to be distinguished. We found two subclusters of intermediate cells and observed that the canonical markers for intermediate cells were all upregulated in only one subcluster (subcluster_0). The upregulation of *Plp1* in this subcluster suggests that it is derived from Schwann cell precursors, and consistent with Bonnamour et al., we identified that the majority of intermediate cells encompassed this subpopulation. The second subcluster we identified was much smaller and showed an upregulation of genes involved in metabolic processes, such as *Idh3B* (isocitrate dehydrogenase (NAD(+)) 3 non-catalytic subunit beta), *Urod* (uroporphyrinogen decarboxylase), *Acot13* (Acyl-CoA Thioesterase 13) and *Rhoa* (Ras Homolog Family Member A). The SV is the energy generator for the inner ear and is therefore a highly metabolically active (reviewed in Thulasiram et al., 2022; reviewed in Yu et al., 2021). Our data suggests that a subset of intermediate cells is specifically dedicated towards maintaining these processes, providing a clearer distinction between the intermediate cell populations and a better understanding of the functional SV.

## LIMITATIONS OF THE STUDY

A limitation of our study was that we were not able to capture endothelial cells or pericytes in our single cell sequencing study, which are components of the BLB. Wnt signaling, particularly Frizzled-4 signaling, in endothelial cells is crucial for regulating vascular development and maintenance in the brain and the retina (Y. Wang et al., 2012) and Frizzled-4 knockout mice experience vascular degeneration in these tissues as well as the inner ear (Xu et al., 2004). In fact, dysregulation of the canonical Wnt/β-catenin pathway is the cause of Norrie disease, a progressive vascular disorder that results in hearing loss (Bryant et al., 2022; Pauzuolyte et al., 2023; Rehm et al., 2002). Since we dissect the whole SV including all the components of the BLB for our *in vitro* cultures, it would be pertinent in future work to gain a molecular perspective on the vascular cells of the SV.

## CONCLUSION

Overall, our *in vitro* studies and single cell RNA sequencing provide a comprehensive understanding of the molecular landscape of the SV. By developing a whole-organ system to culture the neonatal and adult SV, we open the door for more comprehensive studies to investigate the SV in development, disease, and aging. Key genes and pathways revealed through single cell RNA sequencing can be further investigated using this platform to develop biological therapies for SV-associated hearing loss.

## ACKNOWLEDGEMENTS AND FUNDING

This work was supported by the Canadian Institutes of Health Research Doctoral Research Award (CIHR/CGS-D; M.R.T), the Raymond Ng Graduate Award and Harry Barberian Scholarship from the University of Toronto (M.R.T), the Krembil Foundation biomedical research grant (A.D), and the Michael and Sonja Koerner Charitable Foundation (A.D). This work was supported (in part) by the Intramural Research Program of the National Institute on Deafness and Other Communication Disorders (NIH Intramural Research Funds ZIA DC000088 to M.H. of the Auditory Development and Restoration Program and ZIC DC000086 to R.J.M. of the NIDCD/NIDCR Genomics and Computational Biology Core), National Institutes of Health. We thank Dr. Martín Basch for his helpful discussions leading to the generation the P1 single cell RNA sequencing dataset. We also acknowledge the imaging facilities at The Hospital for Sick Children and Sunnybrook Research Institute for the use of and assistance with confocal microscopy. This work utilized the computational resources of the NIH HPC Biowulf cluster. (http://hpc.nih.gov).

